# Periodic neglect after frontoparietal lesions provides causal evidence for rhythmic attention sampling

**DOI:** 10.1101/2022.11.07.515418

**Authors:** Isabel Raposo, Sara M. Szczepanski, Kathleen Haaland, Tor Endestad, Anne-Kristin Solbakk, Robert T. Knight, Randolph F. Helfrich

## Abstract

Contemporary models conceptualize spatial attention as a blinking spotlight that sequentially samples visual space. Hence, behavior fluctuates over time even in states of presumed ‘sustained’ attention. Recent evidence suggested that rhythmic neural activity in the frontoparietal network constitutes the functional basis of rhythmic attentional sampling. However, causal evidence to support this notion remains absent. Using a lateralized spatial attention task, we addressed this issue in patients with focal lesions in the frontoparietal attention network. Our results uncovered that frontoparietal lesions introduce periodic neglect, i.e., temporally-specific behavioral deficits that were aligned with the underlying neural oscillations. Attention-guided perceptual sensitivity was on par with healthy controls during optimal phases but attenuated during the less excitable sub-cycles. Theta-dependent sampling (3 – 8 Hz) was causally dependent on prefrontal cortex, while alpha-band sampling (8 – 14 Hz) emerged from parietal areas. Collectively, our findings reveal that lesion-induced high amplitude, low frequency brain activity is not epiphenomenal, but has immediate behavioral consequences. More generally, these results provide causal evidence for the hypothesis that the functional architecture of attention is inherently rhythmic.

## Introduction

Attention is a key cognitive function to overcome the brain’s limited processing capacities by enhancing behaviorally relevant information^1,2^. A multitude of neuroimaging and lesion studies have demonstrated that the frontoparietal network constitutes the neural basis of attention^3–7^. Previously, spatial attention was conceptualized as a ‘static spotlight’ that remains constant over time ^8^. However, several recent experimental findings have shifted this perspective and suggested that attention operates as a ‘blinking spotlight’ that sequentially samples behaviorally-relevant spatial locations^9^. It remains unaddressed if a blinking spotlight constitutes an active mechanism to distribute limited cognitive resources or whether its discrete nature is the direct consequence of the inherently waxing and waning nature of brain activity. In a series of experiments in humans and non-human primates, it has been demonstrated that attention cycles as a function of the underlying neuronal rhythm (~3 – 12 Hz) of the frontoparietal attention network^9–14^. Performance peaked during phases of enhanced perceptual sensitivity (optimal for information sampling), which are interleaved with suboptimal phases of diminished perceptual sensitivity where attention is shifting to a different location. Albeit mounting correlative evidence, to date there is no causal evidence that demonstrates an unequivocal link between frequency- and spatially-specific rhythmic brain activity and the observed rhythmic modulation of attention.

Over the past decades, a large body of research has systematically characterized spatial attention deficits following focal lesions to the frontoparietal attention network, exemplified by hemispatial neglect, mainly observed in patients with right parietal cortex lesions^15–17^. Spatial neglect is characterized by a failure to attend to and perceive the contralesional hemifield. However, it has long been recognized that focal lesions in the attention network are also detrimental to sustained attention, i.e., deficits in maintaining attention over several seconds to minutes^6,18,19^.The electrophysiological correlates of focal lesions in the attention network have largely been identified for early sensory processing, such as reduced amplitudes of the processing negativity and P300 event-related potential^20,21^. However, it is a well-established clinical finding that focal high amplitude, low frequency rhythmic brain activity as observed on scalp electroencephalography (EEG) is indicative of a lesion^22–25^. To date, no study has investigated the effects of frontoparietal lesions on the fine-grained temporal dynamics of attention at the behavioral and electrophysiological level. Furthermore, it remains unclear if the lesion-induced focal slowing of brain activity has immediate functional consequences^26^.

In the present study, we addressed these unanswered questions by combining whole-head EEG recordings with a well-established task probing attention on the rapid timescale of (sub-cycle) oscillatory brain activity. To determine the causal contributions of distinct nodes of the attention network to rhythmic attentional sampling, we assessed participants with focal lesions in either the prefrontal (PFC) or parietal cortex (PCtx) as well as age-matched healthy controls. We tested whether lesions disrupt the temporal organization of the attention network altering rhythmic sampling behavior. We further assessed if lesions in different network nodes exhibit distinct spectral signatures, which could reflect unique functional contributions to the sequential sampling of the environment, even in states of ‘sustained’ spatial attention. Based on the well-known spatial distribution of brain oscillations ^27,28^, we predicted that prefrontal lesions would impair rhythmic attentional sampling in the theta band (~3 – 8 Hz) ^9,29^, while parietal lesions should disrupt perceptual alpha-band rhythmic sampling (~8 – 14 Hz) ^9,30^.

## Results

We recorded 64-channel electroencephalography (EEG) from patients with chronic focal lesions in either the lateral prefrontal cortex (PFC) or parietal cortex (PCtx) to assess their unique causal contributions to rhythmic attentional sampling. Twenty-five patients with unilateral focal lesions (PFC: n = 13 (6 left/7 right), 57 ± 9 years; PCtx: n = 12 (6 left/6 right, 67 ± 21 years, mean ± SD) and 23 age-matched healthy controls completed a lateralized spatial attention task (**Figure 1A**). Participants were cued to covertly attend either the left or right visual field and responded to a target after a variable cue-target interval (1000 – 2000 ms)^13,31,32^.

**Figure 1.**
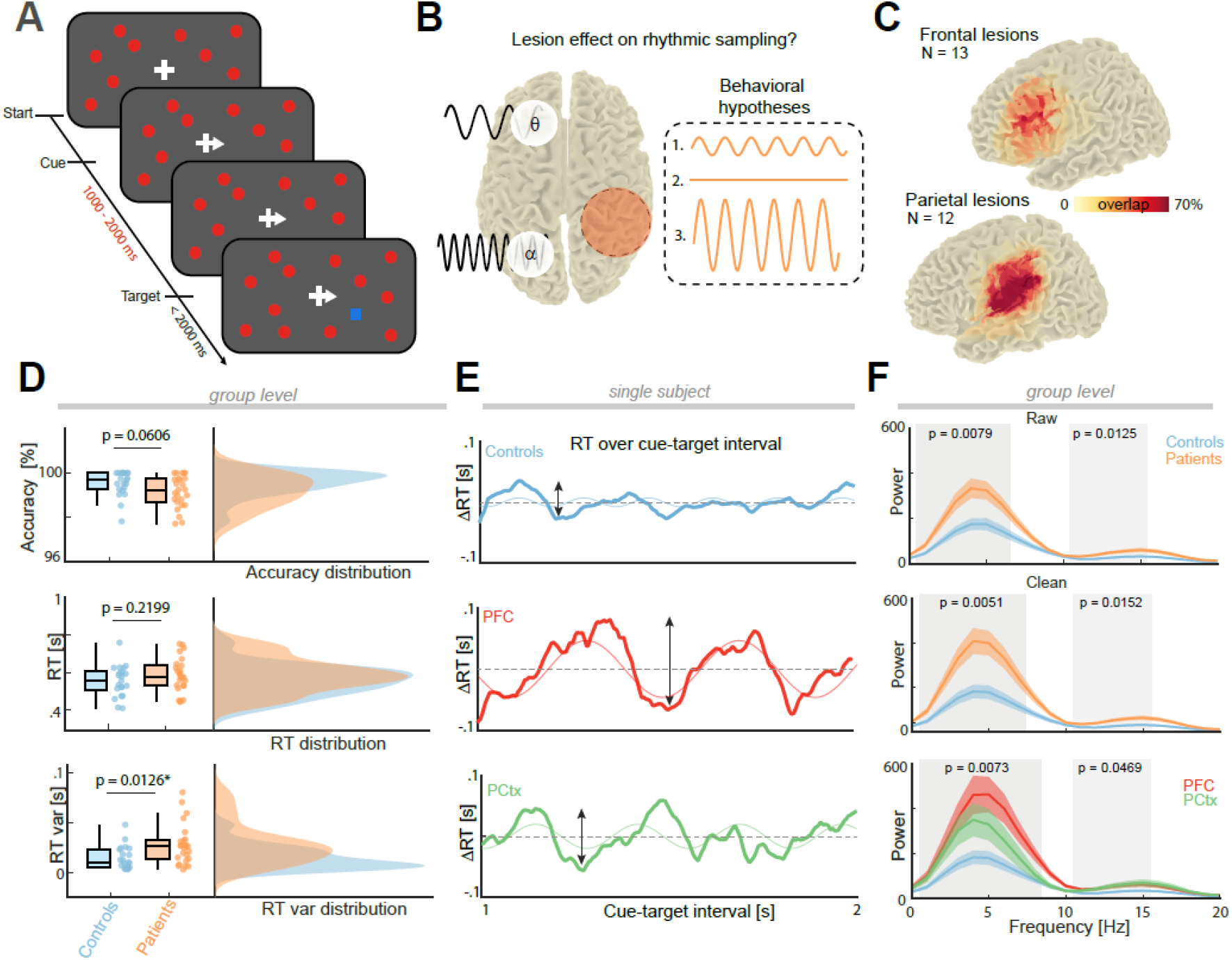
Focal lesions of the frontoparietal network increase rhythmic attentional sampling. **(A)** Illustration of task design. Participants fixated a central fixation cross on a dynamic background with several visual distractors that were randomly switched on or off (red). A centrally presented spatial cue indicated with high probability (70%) the hemifield participants should covertly attend to. After a variable cue-target interval (1000-2000 ms) a target (blue square) appeared, and participants responded with a button press if the target was presented in the cued hemifield. **(B)** Hypothesized task outcomes. Prefrontal cortex (PFC), associated with theta-dependent attention allocation, and parietal cortex (PCtx), linked to the alpha-band activity, comprise the frontoparietal attention network. We predicted that a disruption of the frontoparietal network could lead to one of three possible frequency-specific behavioral scenarios: Attenuated rhythmic behavioral fluctuations (amplitude decrease), suppression of rhythmic sampling or an increased rhythmic sampling (amplitude increase). **(C)** Lesion reconstruction: PFC and PCtx lesion overlapped (in %) for all 25 patients (13 PFC; 12 PCtx) normalized to the left hemisphere. See Figure S1 for single subjects. **(D)** Accuracy, reaction time (RT) and RT variance per group (whiskers indicate maximum and minimum; dots correspond to individual participants). Age-matched controls (blue) and patients (orange) only differed in the reaction time variance, with larger variability in the patients. **(E)** Demeaned, time-resolved RTs and model fit (unconstrained sine wave, thin line) as a function of the cue-target interval for one exemplary participant per group (controls: blue; PFC: red; PCtx: green). Note, RT courses varied periodically across time with overall larger amplitudes in both patients. **(F)** Group-level 1/f-corrected power spectra of the behavioral time courses. Top: Patients exhibited a frequency-specific increase in rhythmic attentional sampling in the theta (2 – 7 Hz; d = −0.80) and high alpha band (12 – 16; d = −0.74). Center: This effect became more pronounced after rigorous exclusion of eye movements (2 – 7 Hz: d = −0.89; 12 – 16 Hz: d = −0.83). Bottom: The increased amplitude in the theta- and alpha-band was also present in both patient groups.

We considered three possible scenarios for how insults to the frontoparietal attention network could impact behavior on a fine-grained temporal scale (Figure 1B): Lesions could disrupt the attention network and either (i) attenuate or (ii) suppress the rhythmic sampling of attention. These scenarios imply that rhythmic sampling constitutes an *active* process to distribute limited cognitive resources across time. However, when assuming that rhythmic behavioral sampling is the direct consequence of underlying rhythmic brain activity (thus, reflecting a *passive* phenomenon), then lesion-induced low frequency activity could (iii) result in an increase in rhythmic attentional sampling.

### Behavioral and Modelling Results: Focal lesions in the frontoparietal network increase rhythmic attentional sampling

Both age-matched controls and patients (**Figures 1C** and **S1A**) performed the lateralized spatial attention task with high accuracy (**Figure 1D** top; controls: 99.53% ± 0.57%, patients 99.16% ± 0.71%, mean ± SEM; t_44_ = 1.92, p = 0.0606, d = 0.57; two-tailed t-test). Moreover, mean reaction times (RTs) did not differ (**Figure 1D** center; t_44_ = −1.24, p = 0.2199, d = −0.37; controls 555 ± 83 ms, patients 587 ± 90 ms, mean ± SEM), while RT variance was increased in patients (**Figure 1D** bottom; t_44_ = −2.60, p = 0.0126, d = −0.77; controls 14 ± 11 ms, patients 27 ± 20 ms, mean ± SEM). Increased RT variance was present in both PFC and PCtx lesion groups (PFC: t_32_ = −1.82, p = 0.0391, d = −0.64; PCtx t_31_ = −2.89, p = 0.0035, d = −1.05; one-tailed t-test) and did not differ between them (t_23_ = −1.31, p = 0.2041, d = −0.52). This effect was independent of the lesioned hemisphere (**Figure S1B**). The observation of systematically increased RT variance in lesion patients raised the question whether this increase exhibited a consistent temporal structure. To address this, we assessed RTs as a function of the cue-target interval (**Figure 1E**; 50ms moving window in steps of 1ms).

To quantify the frequency and oscillatory power, we spectrally decomposed the behavioral traces using a Fast Fourier Transform (FFT). We removed the non-oscillatory 1/f contribution to obtain a whitened power spectrum. First, we considered all available trials. Cluster-based permutation testing revealed narrow-banded power increases in patients in the theta (**Figure 1F** top; 2 – 7 Hz: p_cluster_ = 0.0079, d = −0.80) and alpha band (12 – 16 Hz: p_cluster_ = 0.0125, d = −0.74). To control for a possible impact of eye movements, the analysis was repeated after excluding all trials that contained eye movements. This control analysis strengthened the initial observation (**Figure 1F** center; note the increase in effect size: 2 – 7 Hz: p_cluster_ = 0.0051, d = −0.89; 12 – 16 Hz: p_cluster_ = 0.0152, d = −0.83). Increased rhythmic attentional theta- and alpha sampling was observed in both patient groups independently (**Figure 1F** bottom; PFC: 2 – 8 Hz: p_cluster_ = 0.0057, d = −1.23; 10 – 16 Hz: p_cluster_ = 0.0057, d = −0.92; PCtx: 2 – 5 Hz: p_cluster_ = 0.0311, d = −0.85; 12 – 16 Hz: p_cluster_ = 0.0199, d = −0.91). Notably, increased power was present regardless of whether the stimuli were presented in the ipsilesional or contralesional hemifield (no significant cluster, all uncorrected p > 0.2655; **Figure S1C**). Lastly, to determine the impact of lesion size, we correlated lesion size (number of voxels) and behavioral power across frequencies. This analysis indicated that lesion size was not correlated with behavioral power (p_cluster_ = 0.1508; cluster-based correlation). Collectively, this set of behavioral findings provides robust evidence for the hypothesis that a lesion in the frontoparietal network increases rhythmic attentional sampling (cf. scenario 3; **Figure 1B**).

### Increased rhythmic attentional sampling in lesion patients is phase-dependent

After having established that patients displayed increased rhythmic behavioral fluctuations in the theta- and alpha-bands, we next sought to determine its precise temporal evolution. We conceived three models based on the behavioral result in the frequency domain to explain the observed spectral pattern in the time domain (Figure 2A): (1) Patients might exhibit a similar mean but an overall increased oscillatory amplitude, implying that patients respond faster or slower than controls, depending on the oscillatory phase. (2) Patients might respond slightly slower (nonsignificant offset), albeit with overall stronger fluctuations. In this scenario, patients would perform to par with controls during optimal phases, but worse at suboptimal time points. (3) When both offset and amplitude increase, then patients should perform worse at both, optimal and suboptimal phases, as compared to healthy controls. To test which model best explained the behavioral results, we compared the worst and best RT of every participant, as well as the mean and the amplitude, in the raw behavioral traces (Figure 2B) as well as isolating the slow fluctuations through band-pass filtering (Figure S2).

**Figure 2.**
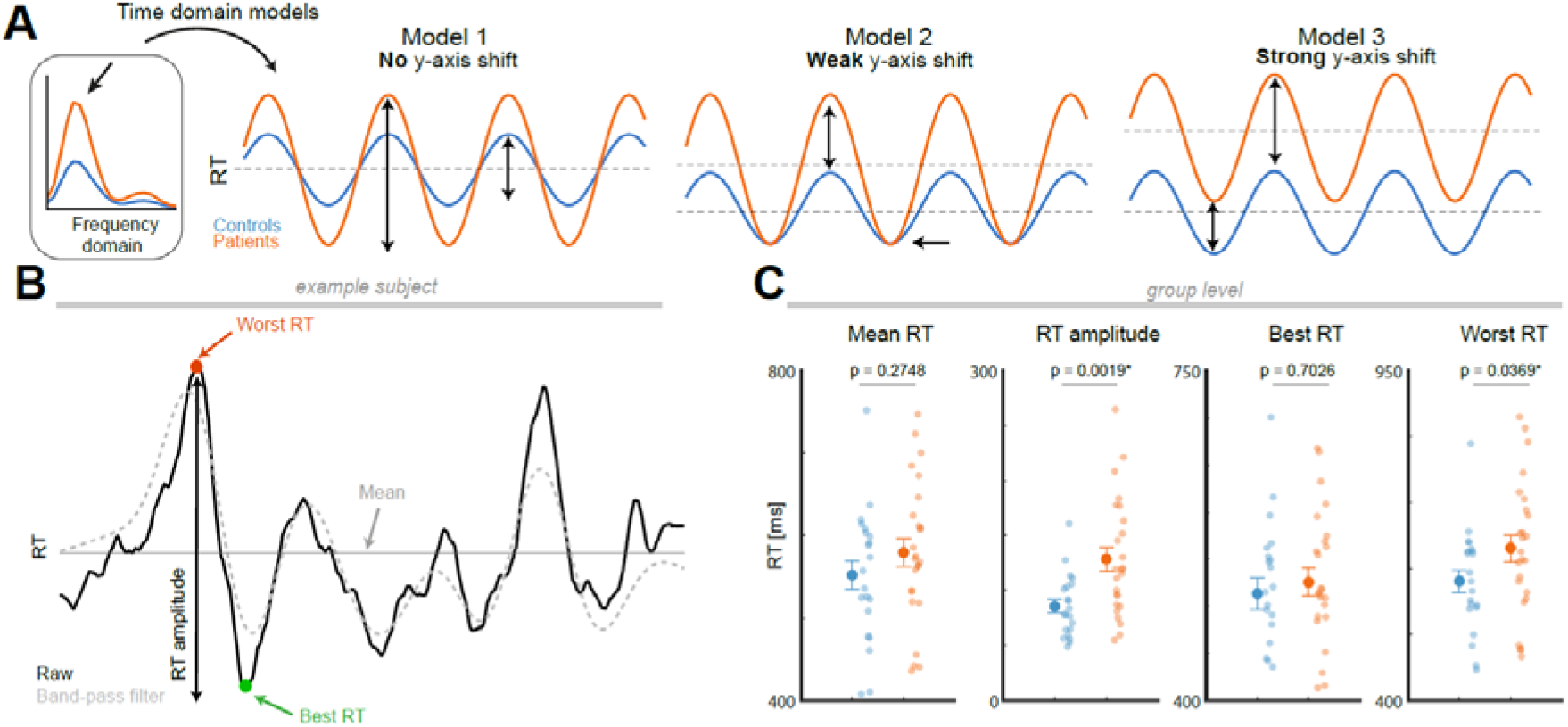
Model comparison of the temporal evolution of behavioral deficits in lesion patients. **(A)** Schematic of hypothesized behavioral models underlying increased rhythmic sampling in patients. Three scenarios were conceivable based on the difference on the frequency domain: Model 1: Comparable mean, but increased variance (amplitude), implying that patients (orange) should exhibit faster RTs at optimal time points and slower RTs during less favorable phases as compared to controls (blue). Model 2: Different mean (non-significant offset along the y-axis) and increased amplitude. Hence, behavioral performance in patients should be on par with controls during optimal phases, but significantly worse during suboptimal phases. Model 3: Different mean, increased amplitude. Hence, performance at optimal and suboptimal phases should be worse in patients. **(B)** Behavioral time-course of one representative patient. Points of interest are defined on the raw (solid line) and band-pass filtered traces (dashed line; **Figure S2**). We defined (i) mean, (ii) amplitude, and (iii) best behavior as well as (iv) worst behavior in the time domain. **(C)** Patients exhibited no significant difference in mean (p = 0.2748) or best RT (p = 0.7026), but RT amplitude (p = 0.0019) and worst RT (p = 0.0369) were significantly increased in patients. Collectively, these observations support model 2 and demonstrate that lesions in the frontoparietal network are detrimental for rhythmic attentional sampling in a phase-specific manner.

To test these models, we first compared mean reaction times, which did not differ (**Figure 2C**; t_44_ = −1.10, p = 0.2748, d = −0.33; two-tailed t-test), replicating the initial results (cf. **Figure 1D**). The overall amplitude of the behavioral time course was increased in patients (t_44_ = −3.30 p = 0.0019, d = −0.98; two-tailed), confirming and extending the observations in the frequency domain (cf. **Figure 1F**). We then tested whether patients were faster at the best phase (the key prediction of model 1) but did not find evidence for this consideration (t_44_ = −0.53, p = 0.7026, d = −0.16; one-tailed). Likewise, we did not find support for systematically slower RTs as predicted by model 3 (t_44_ = −0.53, p = 0.2974, d = −0.16; one-tailed). Finally, our findings favored the prediction that was common to all three models, namely that patient performance is decreased during the worst phase (t_44_ = −1.83 p = 0.0369, d = −0.54; one-tailed). Collectively, these observations support the predictions of model 2. These findings establish temporally-specific behavioral deficits in patients suffering from chronic cortical lesions, since patients performed on par with healthy controls during optimal time points, but significantly worse during suboptimal phases, providing evidence for periodic behavioral neglect.

These behavioral results make several specific predictions regarding the underlying neurophysiology. (1) Given that the patients performed the task with high accuracy, we hypothesized that indices of sensory processing, i.e., early evoked responses, remain largely intact. (2) High amplitude, low frequency EEG activity is indicative of an underlying cortical lesion ^23,24^, which in turn might predict the enhanced amplitude in rhythmic behavioral sampling. (3) Moreover, we observed a clear distinction into best and worst phases in both patients and controls, which implies that the phase-behavior relationships are maintained following a lesion. (4) Lastly, our behavioral results suggest that theta rhythmic sampling is stronger in PFC patients, while alpha band sampling is stronger in PCtx patients (cf. **Figure 1F**); indicating that frontal cortex is the main source driving theta activity, while alpha activity dominates in parietal areas.

### Electrophysiological Results: Focal lesions increase low frequency activity in the frontoparietal network

To assess the neural correlates of increased rhythmic attentional sampling in patients, concomitant EEG was acquired in all participants. After preprocessing and artifact removal, we first assessed cue-locked and target-locked event-related potentials (ERPs). We observed that ERPs were similar in both groups (**Figure 3A**; no significant cluster difference; smallest p_cluster_ = 0.2038). We replicated the previously reported attenuation of early components in PFC lesion patients in the ipsilesional hemisphere ^33^ (**Figure 3B**; t_12_ = 2.69, p = 0.0197, d = 0.45). This effect was not observed for the PCtx group (t_11_ = 0.22, p = 0.8267, d = 0.03) and was temporally-specific (see **Figure S3A** for P300 analysis). Critically, we did not observe any group differences during the behaviorally-relevant cue-target interval (1 – 2 s).

**Figure 3.**
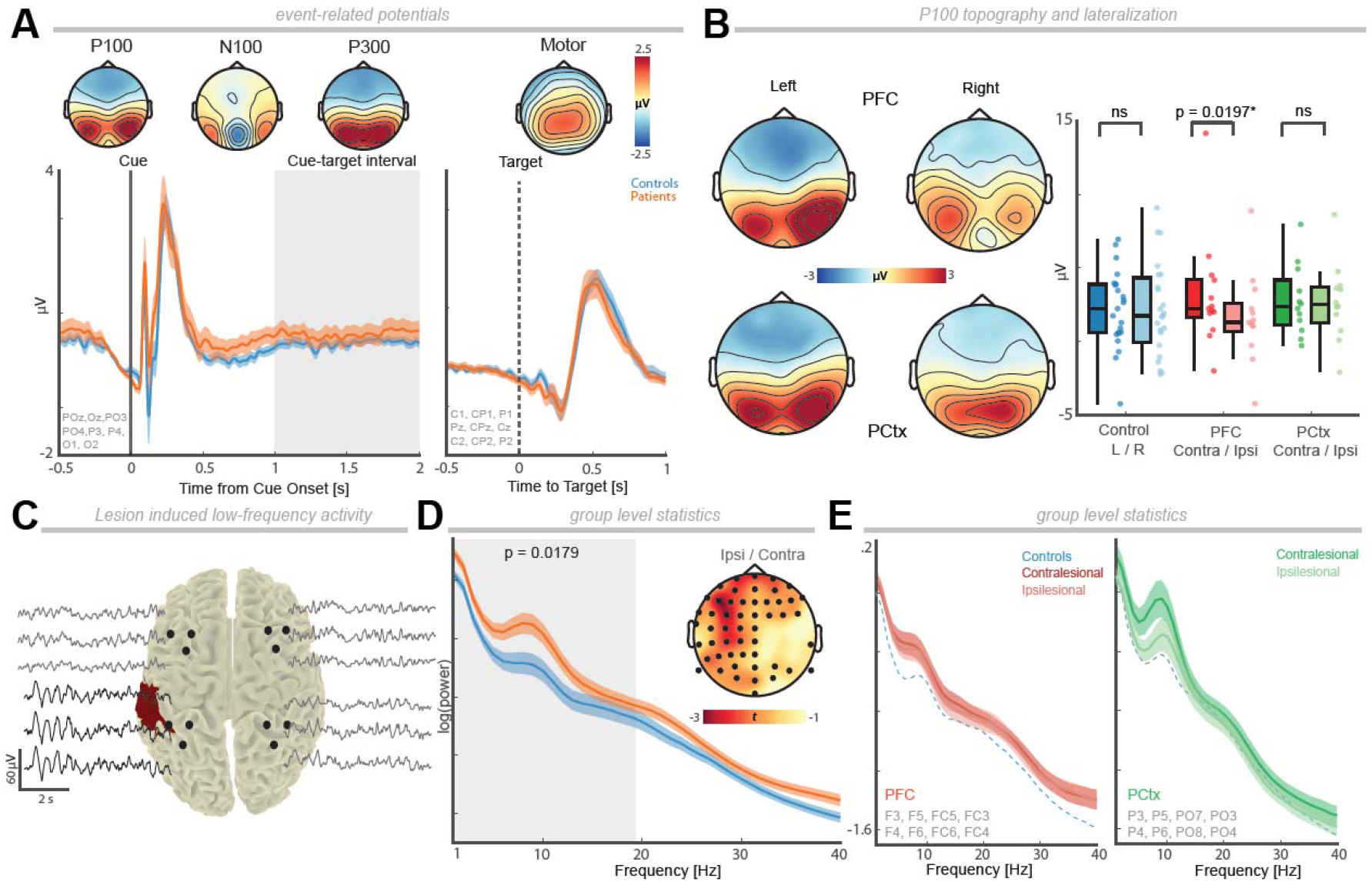
Focal lesions increase low frequency activity in the frontoparietal attention network. **(A)** Grand-average event-related potentials (ERPs; mean ± SEM) cue- (left; posterior channels) and target-locked ERP (right; central channels) demonstrate that ERPs did not differ between patients (orange) and controls (blue). Topographies of the main ERPs (P100, N100, P300, and motor response) averaged across all subjects. (**B**) Left: P100 topography for PFC and PCtx patients with either a lesion in the left or right hemisphere. Note that scalp distributions remained largely intact. Right: Mean P100 ERP (80 – 110 ms) per group (contra-vs ipsilesional posterior channels). Activity over ipsilateral posterior channels was attenuated in PFC lesion patients. **(C)** Illustration of increased, peri-lesional low-frequency EEG activity. **(D)** Grand-average power spectra with mirrored electrodes in right hemisphere lesion patients, so that lesions are all located over the left hemisphere (mean ± SEM; black dots indicate significant channels; **Figure S3** for non-mirrored electrode positioning). Patients exhibited widespread increased low frequency power in comparison to controls (1-19 Hz; p_cluster_ = 0.0179). **(E)** Left: Grand-average power spectra of contra- and ipsilesional PFC channels, revealing no significant differences. The blue dashed line highlights the mean of the control group. Right: PCtx group. Same conventions as in the left panel. Again, no significant difference was observed between ipsi- and contralesional channels for PCtx lesion patients.

We observed a well-known clinical finding, namely that channels over the lesioned tissue display high-amplitude, low-frequency activity (**Figure 3C**). To quantify this observation, we spectrally decomposed the electrophysiological time series during the cue-target-interval by means of a Fourier transform. We observed increased power in the low frequency range (1 – 16 Hz) across the majority of EEG sensors in patients as compared to healthy controls (**Figure S3B** left; p_cluster_ = 0.0259, d = −0.58). To determine whether this power increase was pronounced over the lesioned hemisphere, we mirrored all the electrodes across the midline in patients with right hemisphere lesions, which again revealed a large bihemispheric cluster (**Figure 3D**; 1 – 19 Hz; p_cluster_ = 0.0179, d = −0.55). To further determine the impact of a focal lesion on brain-wide spectral power, we re-referenced the EEG signal to a unipolar reference that was not overlaying the lesion (Cz). This again replicated a wide-spread increase in lower frequency activity in patients (**Figure S3B** right; p_cluster_ = 0.0149, d = −0.49). Finally, we compared activity at ipsi- and contralesional electrodes. Both PFC and PCtx lesions resulted in a comparable, widespread power increase (**Figure 3E**; no significant cluster was found for these comparisons, all uncorrected p > 0.0965). In sum, these findings demonstrate a wide-spread increase of low frequency EEG activity in lesion patients.

### Theta and alpha oscillations predict increased rhythmic attentional sampling

After having observed a systematic increase in low frequency activity in both behavior (**Figure 1F**) and electrophysiology (**Figure 3D**) in patients, we next tested if these increments were directly correlated. In the control group, we first established that mean behavioral and EEG power were significantly positively correlated across multiple lower frequencies with two distinct peaks at 4 Hz and 11 Hz (**Figure 4A** left; 1 to 20 Hz: r = 0.507, p_cluster_ = 0.0110; cluster-corrected correlations); thus indicating that these phenomena were not independent. To further quantify the relationship between behavioral and EEG activity in patients, we introduced a composite metric to quantify their mutual dependence for every subject, channel, and frequency, termed the rhythmic sampling index (RSI). The RSI was defined as the resultant vector length in a 2D space spanned by frequency-specific behavior and EEG power (**Figure 4A** right). Patients exhibited a larger RSI in comparison to controls in the lower-frequency range (**Figure 4B** left; 1 – 13 Hz: p_cluster_ < 0.0001, d = −0.7283), indicating that patients’ larger neural oscillation’s amplitude predict larger behavioral oscillatory power (**Figure 4B** right).

**Figure 4.**
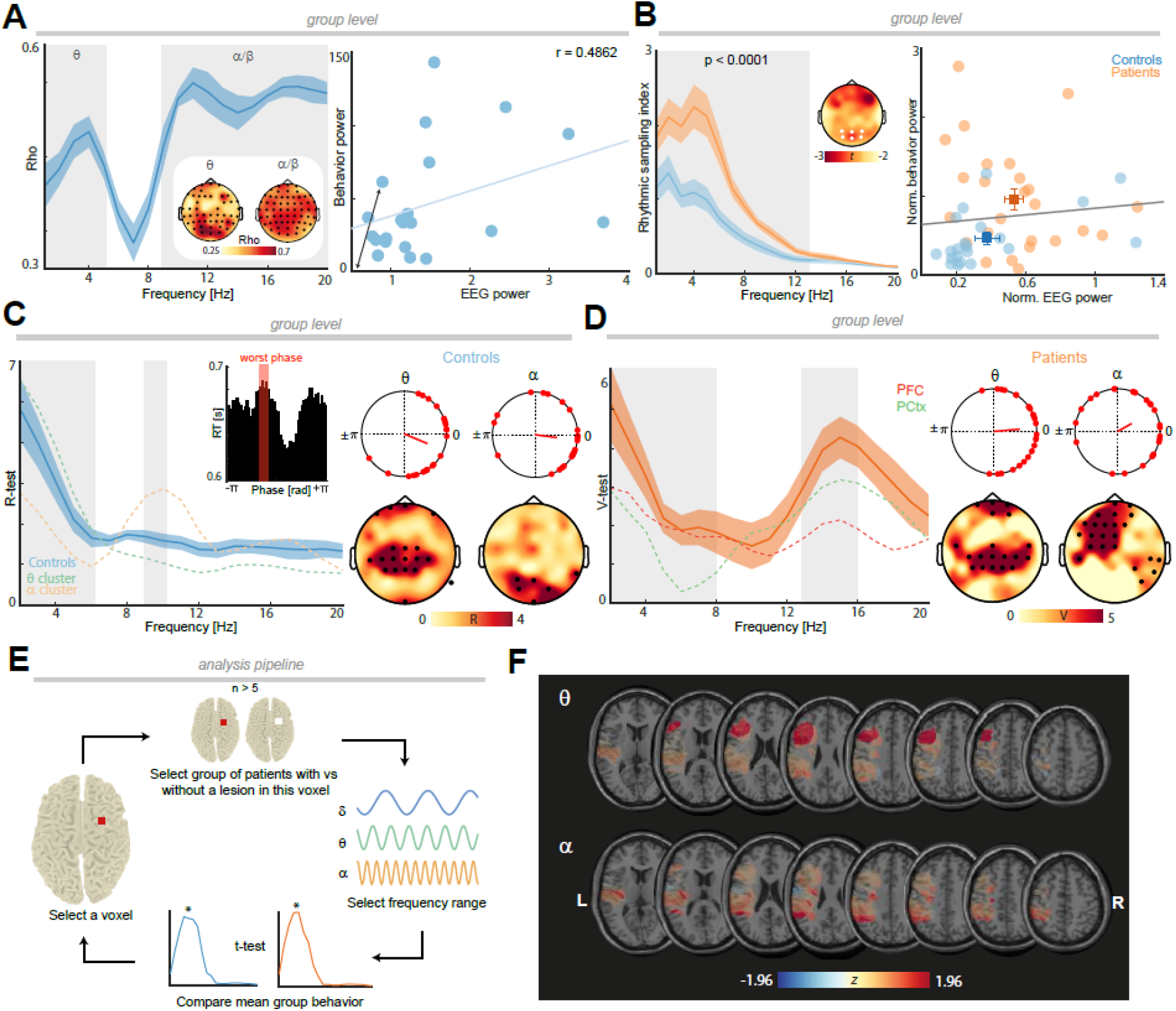
Neuronal oscillations predict rhythmic attentional sampling. **(A)** Left: Correlation of spectrally-resolved behavior and electrophysiology in healthy controls (cluster-corrected correlation: p_cluster_ = 0.0110; shaded error bars indicate bootstrapped correlation coefficient standard error across 100 repetitions) revealed a large cluster spanning multiple channels and frequencies with two distinct peaks in the theta and alpha/beta range (gray shaded areas; thresholded at rho = 0.3). Inset: Topographies indicate the spatial extent (black dots indicate cluster electrodes). Right: Neuronal oscillatory power of theta cluster as a function of behavioral power (averaged across all significant theta cluster channels; the blue line highlights the linear regression). To quantify the relationship of EEG and behavioral power, we calculated the rhythmic sampling index (i.e., the magnitude of the black arrow after normalization). **(B)** Left: Patients had an increased association between behavioral and EEG power in lower frequencies (1 – 13 Hz; p < 0.0001). Right: Normalized EEG power as a function of normalized behavioral power for healthy participants and patients in the six posterior example electrodes illustrated in the topography (squares, mean; error bars, SEM). **(C)** Left: Phase-resolved behavior as a function of frequency in healthy controls to assess if the phase that was associated with worst performance (cf. **Figure 2A**, *model 2)* was consistent at the group level. The inset demonstrates the phase-resolved reaction times in the theta range (single subject and electrode example). Note the non-uniform distribution across the phase bins (worst phase in red). Rayleigh tests identified consistent phase clustering in the theta (2 – 6 Hz) and alpha (9 – 10 Hz) range (gray shaded areas depict significant frequencies, FDR-corrected p < 0.05; shaded error bars indicate SEM across channels; dashed lines indicate the within-cluster average; cf. right panels). Right: Spatial extent of the phase clustering in the theta- and alpha-bands (black dots indicate significant electrodes). Significant theta phase consistency was observed over fronto-centro-parietal sensors, while alpha phase consistency was observed over occipital sensors. **(D)** Left: Phase-behavior relationships remained stable in lesion patients. We observed a highly comparable mean direction (V-test against known mean direction in healthy controls; cf. panel C) in patients in the theta (2 – 8 Hz) and high alpha band (13 – 16 Hz; shaded gray areas correspond to significant frequencies; FDR corrected p < 0.05; shaded error bars indicate SEM across channels; dashed lines represent the mean test statistic V of PFC and PCtx patients). Significant phase clustering in the same direction as healthy controls indicates that the non-uniform relationship between phase and behavior persisted after focal lesions. Right: Spatial extent of the phase clustering. Again, theta phase clustering emerged mainly over fronto-central sensors, while alpha phase clustering was apparent over centro-parietal sensors. **(E)** Illustration of the spectrally-resolved voxel-based lesion-symptom mapping (VLSM) analysis. For every voxel, patients were divided into two groups depending on whether this voxel was within the lesion or not. Next, frequency-specific behavioral power (1 – 20 Hz; cf. **Figure 1F**) was compared by a *t*-test. The analysis was then repeated on a voxel-by-voxel basis. T-values were transformed into z-scores. P-values across all voxels were FDR-corrected. **(F)** VLSM maps depict *z*-values of all (p < 0.05; uncorrected) voxels in the theta (2 – 7 Hz) and alpha (12 – 16 Hz range; frequency range analogous to **Figure 1F**). See also **Figure S4** for FDR-corrected maps. Lesions within the lateral prefrontal cortex predicted a behavioral theta power increase, whereas parietal deficits predicted an increase in rhythmic sampling in the alpha band.

### Phase dependence of rhythmic attentional sampling is preserved in patients

Having established that a lesion-induced increase of low frequency EEG activity predicts an increase in rhythmic attentional sampling in behavior, we next tested a key prediction of model 2 (**Figure 2A**), which implied that precise oscillatory phase that governs behavior should be consistent in controls and patients. Specifically, we determined the suboptimal phase where the slowest reaction time occurred. To quantify the association between oscillatory phase and behavior, we computed phase-resolved reaction times (**Figure 4C** left, inset). We divided the phase into 50 equally distributed bins (ranging from–π to +π) and computed the average reaction time for all trials within a 90° window centered on every phase bin. The suboptimal phase (slowest reaction time) was then determined for every subject per channel and frequency. To test for phase clustering per frequency, we first conducted Rayleigh tests in healthy controls for each channel and frequency, separately. We observed significant phase clustering in the theta range (**Figure 4C**; 2 – 6 Hz; −24.9° ± 10.1°, circular mean ± SEM; resultant vector length r = 0.64, Rayleigh z = 3.21, FDR corrected all p ≤ 0.0075) and alpha band (9 – 10 Hz; −7.4° ± 11.8°, circular mean ± SEM; resultant vector length r = 0.51, Rayleigh z = 2.16, FDR corrected all p ≤ 0.0215), indicating that the phase of theta and alpha activity predicts behavior. Next, we assessed whether patients exhibited the same preferred phase as controls (nonuniform phase distribution around the mean phase in healthy controls; V-test). We observed significantly similar phase clustering in the theta (**Figure 4D**; 2 – 8 Hz; 5.2° ± 10.2°, circular mean ± SEM, v = 3.08, FDR corrected all p ≤ 0.0075) and alpha bands (13 – 16 Hz; 29.4 ± 12.8°, circular mean ± SEM, v = 4.33, FDR corrected all p ≤ 0.0059), thus suggesting that phase-behavior relationships in patients remained intact. Furthermore, these results establish that a clear separation into optimal and suboptimal phases for behavior was maintained after focal lesions, despite the overall amplitude increase. Taken together, these findings reveal that the increased rhythmic attentional sampling is not a consequence of altered phase-behavior dependencies but is a direct result of increased low frequency power.

### Dissociable neural origins of theta- and alpha-band rhythmic sampling

Lastly, we determined how different nodes of the frontoparietal network contributed to the theta and alpha-band rhythmic attentional sampling. We employed spectrally-resolved voxel-based lesion symptom mapping (VLSM) (Figure 4E/F) to assess the unique contribution of every voxel to frequency-specific rhythmic behavior. We observed that the theta behavioral cluster was associated with lesions in the lateral prefrontal cortex (Figure 4F, thresholded at z = ±1.96; 2 – 8 Hz; d = 1.00, see Figure S4 for FDR-corrected maps), while the late alpha behavioral cluster was associated with lesions in the temporoparietal junction (12 – 16 Hz; d = 1.11). Collectively, spectrally-resolved VLSM provides causal evidence for the hypothesis that theta-dependent rhythmic attentional sampling originates in the PFC, while alpha-band sampling originates from parietal regions.

## Discussion

Our results demonstrate that chronic lesions in the human frontoparietal attention network cause periodic neglect, where patients exhibit temporally-specific behavioral deficits on the rapid timescale of oscillatory brain activity. Critically, the current findings revealed that lesion-induced high amplitude, low frequency brain activity is not epiphenomenal, but has immediate functional consequences on attention. While patients performed on par with controls during optimal phases, attention allocation was attenuated during suboptimal time windows. These results provide causal evidence for the hypothesis that low frequency oscillations provide the functional substrate for rhythmic sampling of the environment^2^ and more broadly, reveal their causal role for the rhythmic nature of cognition^34^.

### A rhythmic theory of attention

Classic theories conceptualized attention as a ‘static spotlight’ that prioritizes perception at an attended location ^8^ or object ^35^ and has its neural basis in the frontoparietal network ^3^. In these models, attention is assumed to be constant over time to continuously boost sensory representations. However, several recent findings are incompatible with the traditional notion of a ‘static spotlight’^10,13,36,37^. When behavior was probed on a fine-grained temporal scale, spatial attention was shown to fluctuate over time at a theta rhythm (3 – 8 Hz)^14^. Recently, similar observations have been made for object-^10^ and feature-based ^38^ attention as well as abstract cue-guided visual perception^12^. It might also apply to related cognitive concepts, such as working-memory^39^ or other sensory domains, such as audition^40^. Several lines of inquiry suggested that rhythmic brain activity in the frontoparietal attention network constitutes a viable mechanism that efficiently segregates attentional sampling from attentional shifting, i.e., separates sensory from motor functions ^2^. Specifically, electrophysiological recordings in non-human primates^11,41^ and humans^13,42^ demonstrated that theta oscillations shape neural excitability in a phase-specific manner and periodically reweight functional connections between different nodes in the frontoparietal attention network^11,42,43^. Recently, the rhythmic nature of attention has been called into question based on behavioral modeling^44^ and is now actively debated^45^. These controversial findings are a direct consequence of the lack of causal data linking rhythmic fluctuations in behavior to oscillatory brain activity.

Here, we provide causal evidence based on the systematic study of patients with focal lesions in key nodes of the attention network, i.e., the prefrontal and parietal cortex. Our results revealed increased rhythmic attentional sampling in lesion patients. While both PFC and PCtx patients exhibited increased sampling in the theta- and alpha-bands, VLSM localized theta-band sampling to the PFC and alpha-band sampling to the parietal cortex, in line with their presumed neural origins^43,46^. On the functional level, theta- and alpha-band activity predicted rhythmic sampling behavior in a frequency-specific manner. This link was already apparent in the intact brain (**Figure 4A**), but markedly increased following a lesion given the pronounced increase of low frequency band activity (**Figure 3D** and **4B**). Collectively, enhanced rhythmic sampling was driven by the increased amplitude, while the oscillatory organization into excitatory and inhibitory sub-cycles was preserved following focal lesions (**Figure 4D)**. In sum, this set of findings provides causal evidence for the contention that the inherent rhythmic nature of brain activity constitutes the functional basis of rhythmic attentional sampling.

### Attention deficits in space and time after focal brain lesions

Attention deficits upon focal lesions have also mainly been studied in the spatial domain and are best exemplified by the hemispatial neglect syndrome, which is characterized by reduced awareness of the contralesional side following unilateral brain damage^15,47^. Typically, spatial neglect is caused by inferior parietal lesions in the right hemisphere^48^, affecting mostly the contralesional visual field. However, spatial neglect can also be observed after cortical or subcortical lesions^16,49,50^ and occasionally also affects the ipsilesional visual field^47,51^. Neglect is often pronounced in the (semi-) acute phase, where patients exhibit increased RTs to stimuli presented in the contralesional hemifield^31^. Neglect has been shown to be time-dependent, albeit on longer timescales than reported here. For example, the attentional blink^52^ was markedly prolonged to ~1400ms (as compared to ~400 ms) in neglect patients^53^, even when stimuli were presented centrally. Likewise, neglect patients exhibit aberrant inhibition-of-return, as they show facilitation rather than inhibition for repeated events on the non-neglected hemifield^54,55^. While time-dependent attention deficits over several seconds have been described in neglect patients^52–54^, no work has examined attention deficits on the fine-grained temporal scale of brain oscillations.

In the present study, chronic stroke patients were assessed who did not exhibit spatial neglect per se (**Figure S1B**). However, behavioral deficits were evident as increased response time variability, in line with previous reports in frontal lesion patients^56,57^, which had not been linked to electrophysiological brain activity. Critically, the increased variability exhibited a clear temporal structure with distinct spectral peaks in the theta- and alpha-bands when probed on a fine-grained temporal scale. These frequency- and phase-specific deficits were defined by the lesion-mediated, increased low frequency EEG amplitude. In contrast to spatial neglect, no lateralization was observed (**Figure S1C**).

Collectively, these observations reveal a ‘periodic neglect’ with reduced awareness during specific moments in time, which directly correspond to time windows as defined by the inhibitory sub-cycle of low frequency oscillations. As in spatial neglect, our results demonstrate that deficits emerge after lesions to multiple network nodes, hence, conceptualizing periodic neglect as a network disorder. More broadly, this set of findings posits that rhythmic attention as the functional outcome from rhythmic interactions in the frontoparietal network.

### Brain lesions and EEG slowing

Focal EEG slowing is a well-established clinical hallmark that indexes underlying brain lesions^22–25^, but the neural mechanisms that give rise to this phenomenon remain unclear. While often regarded as physical distortions caused by the damaged tissue ^58^, more recent evidence suggests that low frequency activity may reflect functional reorganization after stroke^23,26^. The emergence of coherent neural activity might index reorganization and has been shown to predict recovery of motor control^26^. However, how these findings translate to higher cognitive functions outside of the motor domain remains unknown. Here, we replicated the well-known effect of increased low frequency amplitude following a focal lesion (**Figure 3D**). Our findings demonstrate that increased power is not limited to the peri-lesional cortex but is widespread across the entire frontoparietal network (**Figure S3B/C**). Critically, the increase in low frequency power also predicted increased behavioral power, i.e., temporally-structured response time variability in a phase-specific manner: During optimal phases, patients performed on par with controls, while behavior was periodically impaired during suboptimal phases. Altogether, these findings establish that coherent, lesion-mediated low frequency activity has an immediate behavioral impact and does not constitute an epiphenomenon. A testable hypothesis for future studies is whether the emergence of low frequency activity is a suitable biomarker to track cognitive recovery, similar to previous findings that implicated low frequency activity in neural plasticity underlying motor recovery after stroke ^23,26^.

### Conclusions

In summary, our results reveal a hitherto unknown behavioral deficit resulting from focal brain lesions to the frontoparietal attention network. We demonstrate that lesion-mediated, coherent low frequency activity introduces a periodic behavioral neglect with reduced awareness for specific moments in time. Specifically, neglected time windows are defined as a less excitable sub-cycle of the underlying neural oscillation. More broadly, these results provide causal evidence for the hypothesis that rhythmic attentional sampling has its neural basis in synchronized frontoparietal network activity ^36,43^. In the future, these insights provide the opportunity to tailor targeted interventions to the phase and frequency of oscillatory brain activity ^59^.

## Acknowledgements

This work was funded by the Baden Wuerttemberg Foundation (Postdoc Fellowship; RFH), German Research Foundation, Emmy Noether Program (DFG HE8329/2-1; RFH) and the Hertie Foundation, Network for Excellence in Clinical Neuroscience (RFH), the Research Council of Norway (grant number 240389; AKS, TE), the Research Council of Norway (Centre of Excellence scheme, grant number 262762; RITMO, RITPART International Partnerships for RITMO Centre of Excellence, grant number 274996; AKS, TE, RTK), Department of Veterans Affairs Research Career Scientist Program (KH) and by an NIMH Conte Center Grant (1 PO MH109429, RTK) and NINDS (2 R01 NS021135, RTK).

## Materials and Methods

### Participants

23 healthy older adults (12 males; mean ± SD [range]: 61 ± 14 [21 – 78] years of age, 17 ± 1.5 years of education), 13 patients with lesions in the lateral prefrontal cortex (6 males; 57 ± 9 [41 – 73] years of age; 16 ± 2.5 years of education) and 12 patients with parietal lesions (6 males; 67 ± 21 [20 – 89) years of age, 15 ± 2.6 years of education) were recruited for this study. Lesions were unilateral (PFC: n = 6 left, 7 right hemisphere; PCtx: n = 6 left, 6 right hemisphere). All lesions were chronic (10.43 ± 7.38 [0.74 - 26] years elapsed) and caused by a single stroke or surgical resection of a low-grade tumor. No evidence of tumor regrowth was detected in any of the tumor patients at the time of testing. Patients were recruited from three different sites. 11 patients were recruited at the University of California, Berkeley, 10 patients were tested at the University of New Mexico’s Health Sciences Center, and 3 patients were tested at Oslo University Hospital. Age-matched controls were recruited at the University of California, Berkeley. Two control participants were excluded from the analyses given insufficient EEG data quality (n=1) and excessive drowsiness (n=1). All subjects had normal/corrected-to-normal vision. The patients were evaluated by a clinician prior to testing and had no other neurological or psychiatric diagnoses. All subjects gave informed consent and all procedures were approved by the Institutional Review Board as well as by the Committee for Protection of Human Subjects at the University of California, Berkeley (Protocol number: 2010-02-783) or the Regional Committee for Medical and Healthy Research Ethics and conducted in agreement with the Declaration of Helsinki.

### Lesion reconstruction

Lesion reconstructions were obtained by manual delineation based on structural MRIs obtained after study inclusion under the supervision of a neurologist. Fluid Attenuated Inversion Recovery (FLAIR), T1 and T2 weighted images of each patient’s brain were co-registered to a T1 MNI Template using Statistical Parametric Mapping software’s (SPM) New Unified Segmentation routine ^62^. Lesion delineation was then performed on axial mosaics of the normalized T1 scans using MRIcron ^65^. The resulting lesion masks were then converted to three-dimensional MNI space using the Mosaic to Volume routine in SPM.

### Behavioral Task

Stimulus presentation was controlled with EPrime software (Psychology Software Tools). Participants sat ~60 cm away from the screen. They performed a spatial attention reaction time task where they had to maintain fixation on a cross on a dynamic background with several visual distractors, which were randomly switched on or off. Participants were cued to either the left or right hemifield by a centrally presented cue (70% validity) and asked to covertly shift their attention to the cued hemifield. After a variable cue-target interval (1000 – 2000ms), a blue square target was presented. Participants were instructed to respond to targets presented in the cued hemifield as quickly and accurately as possible and to withhold a response to targets presented in the opposite hemifield. Participants performed the total duration of the task, consisting of 420 trials.

### EEG and eye position data acquisition

EEG data were collected using a 64 channel BioSemi ActiveTwo with active electrodes mounted on an elastic cap according to the International 10-20 System (BioSemi, Amsterdam, Netherlands), sampled at 1024 Hz. Vertical and horizontal eye movements were monitored by 3 electrodes. Continuous gaze position was additionally recorded to exclude any trials post hoc where eye movements occurred. Berkeley eyetracking data were collected using an Eyelink 1000 optical tracker (SR Research, Ontario, Canada), sampled at 1 kHz. No eyetracking was performed in New Mexico and Oslo.

### Behavioral data analysis

We calculated mean target detection accuracy, mean target reaction time and reaction time variance per group. Stimuli were lateralized during presentation, so we further divided patients depending on their lesion location to test for effects of laterality. Spectral analysis on behavioral time courses was performed on the 1000 – 2000 ms cue-target interval after the cue event. Trials where eye movements occurred were excluded and only correct responses to targets at the cued location were considered. To extract the behavioral time-course, we shifted a 50ms window in steps of 1ms from 1000 – 2000ms and re-calculated the reaction times across all validly cued trials in the respective time window. The traces were smoothed with a 25-point boxcar function, demeaned and linearly detrended. We obtained spectral estimates from a Fast Fourier Transform (FFT) in steps of 1 Hz from 1 – 20 Hz after applying discrete prolate spheroidal sequences (dpss) multi-taper with ±3 Hz spectral smoothing. We attenuated the 1/f background activity by multiplying spectral estimates per frequency-of-interest with the respective center frequency.

### EEG data analysis

Preprocessing: The data were offline re-referenced to a common average, demeaned and linearly detrended, high-pass filtered at 0.3 Hz and low-pass filtered at 70 Hz using finite impulse response filters and re-sampled to 1024 Hz. Line noise harmonics (60 Hz for US data and 50 Hz for Oslo data) were removed using a bandstop filter. The data were then visually inspected for artifacts. Eye movements and excessively noisy epochs and channels were rejected. Channels exhibiting increased noise were then reconstructed by interpolation of the mean of the nearest neighboring channels. Next, the data were submitted to an independent component analysis. We excluded components that resembled muscle, heartbeat, or eye movement artifacts from the remaining channels. The final dataset included an average of 356 trials per subject (± SD [range] trials: ± 37 [266 – 409]. Finally, the data were epoched into 5 s long segments, starting 1 s before trial onset.

Event-related potentials (ERP): We extracted the ERPs from the epoched data after applying an absolute baseline correction (−0.2 to 0 s before cue onset). The EEG data segments were low-pass filtered at 40 Hz and smoothed with a 30-point boxcar function for display purposes. All correct trials were included.

Spectral analysis: Power spectra were obtained during the cue-target interval. Spectral estimates were computed by means of a FFT after applying a Hanning window (1-40 Hz, 1 Hz steps) and zero padding.

EEG-behavior correlation: Behavior and EEG correlation was computed using a cluster-based correlation between the EEG and the behavior power spectrum from 1 to 20 Hz. Rhythmic behavioral sampling was first averaged within the significant frequency bands (2 – 7 Hz and 12 – 16 Hz) per participant to obtain a single value reflecting behavioral rhythmic sampling. This approach was viable, because rhythmic sampling in the theta and alpha bands were not independent (rho = 0.6940, p = 0.0002). Second, we introduced a composite metric to quantify the dependence of EEG and behavioral power termed rhythmic sampling index (RSI). The index was defined as the vector length (Euclidean distance to the origin of the coordinate system) for every subject, channel, and frequency in a 2D space consisting of normalized (divided by the maximum value) behavior and EEG power. Normalization was necessary to equate the differences in absolute values between the behavioral (ms) and EEG (μV^2^) scale.

Phase-behavior correlation: To extract the instantaneous analytic phase, we downsampled the data to 256 Hz and band-pass filtered the data from 2 – 30 Hz (± center-frequency / 4) per frequency band and applied a Hilbert transform to extract the instantaneous phase at target onset. Only trials where the target was successfully detected were included in the analysis. Next, we binned the phase angles at target onset into 50 equally distributed bins and computed the average phase-resolved reaction times per channel and frequency bin across all trials within a 90° window centered around every phase bin. Subsequently, we determined the phase bin with the slowest RT per participant, channel, and frequency for statistical testing.

### Spectrally-resolved voxel-based lesion symptom mapping

Data were further analyzed using an adaptation of voxel-based lesion symptom mapping^66^, which was spectrally-resolved. This method maps the relationship between behavior and brain lesions on a voxel-by-voxel basis. Here, we normalized lesion reconstructions in MNI space for every patient along with their spectrally-resolved behavioral data. The analysis was carried out across all frequencies and subsequently visualized for the behaviorally-relevant frequency ranges (2 – 7 Hz and 12 – 16 Hz) after correction for multiple comparisons across all frequencies and voxels. All lesion maps were flipped onto the left hemisphere to increase statistical power, since we did not observe any differences between lesion hemispheres in all previous analyses. Then, we conducted a *t*-test at every voxel to compare between behavioral power of patients with and without a lesion in that voxel. This approach indexed the brain areas whose damage had the greatest impact on the behavioral power increase in the significant frequency bands. Tests were confined to voxels where there were more than five patients per sub-group (i.e., with and without a lesion). We then z-scored the t-values.

### Statistical testing

Throughout, we report single subject data and highlight effects that generalize across the population and were observed in every participant. Unless stated otherwise, we employed two-tailed paired t-tests (**Figure 1D, 2C** and **3B**) to infer significance at the group level. For the electrophysiological data, we employed cluster-based permutation tests to correct for multiple comparisons as implemented in Fieldtrip (Monte Carlo method; 1000 iterations; maxsum criterion ^67^) based on either paired or unpaired two-tailed t-tests, unless stated otherwise. Clusters were either formed in time (e.g., **Figure 3A**) or in the frequency domain (e.g., **Figure 1F, 3D** and **4B**). We furthermore used cluster-based correlation based on Pearson correlation coefficient, which was subsequently transformed into a t-statistic (e.g., **Figure 4A**). We included bootstrapped standard errors where applicable. In several instances where cluster testing was not feasible (e.g., for circular data or voxel-based lesion symptom mapping), we also employed FDR correction (Benjamini-Yekutieli; q = 0.1). Circular statistics as the Rayleigh test and V-test (**Figures 4C/4D**), which test for circular non-uniformity and non-uniformity with a specified mean direction respectively, were carried out using the CircStat toolbox ^61^. In cases were multiple p-values were obtained (circular data across different dimensions; e.g., frequency and electrodes), we combined p-values using the method by Stouffer et al. to infer significance ^68^ as outlined in detail by VanRullen, 2016a ^64^. Effect sizes were calculated using Cohen’s d, the correlation coefficient rho, or the resultant vector length.

## Supplemental information

**Figure S1 related to Figure 1.**
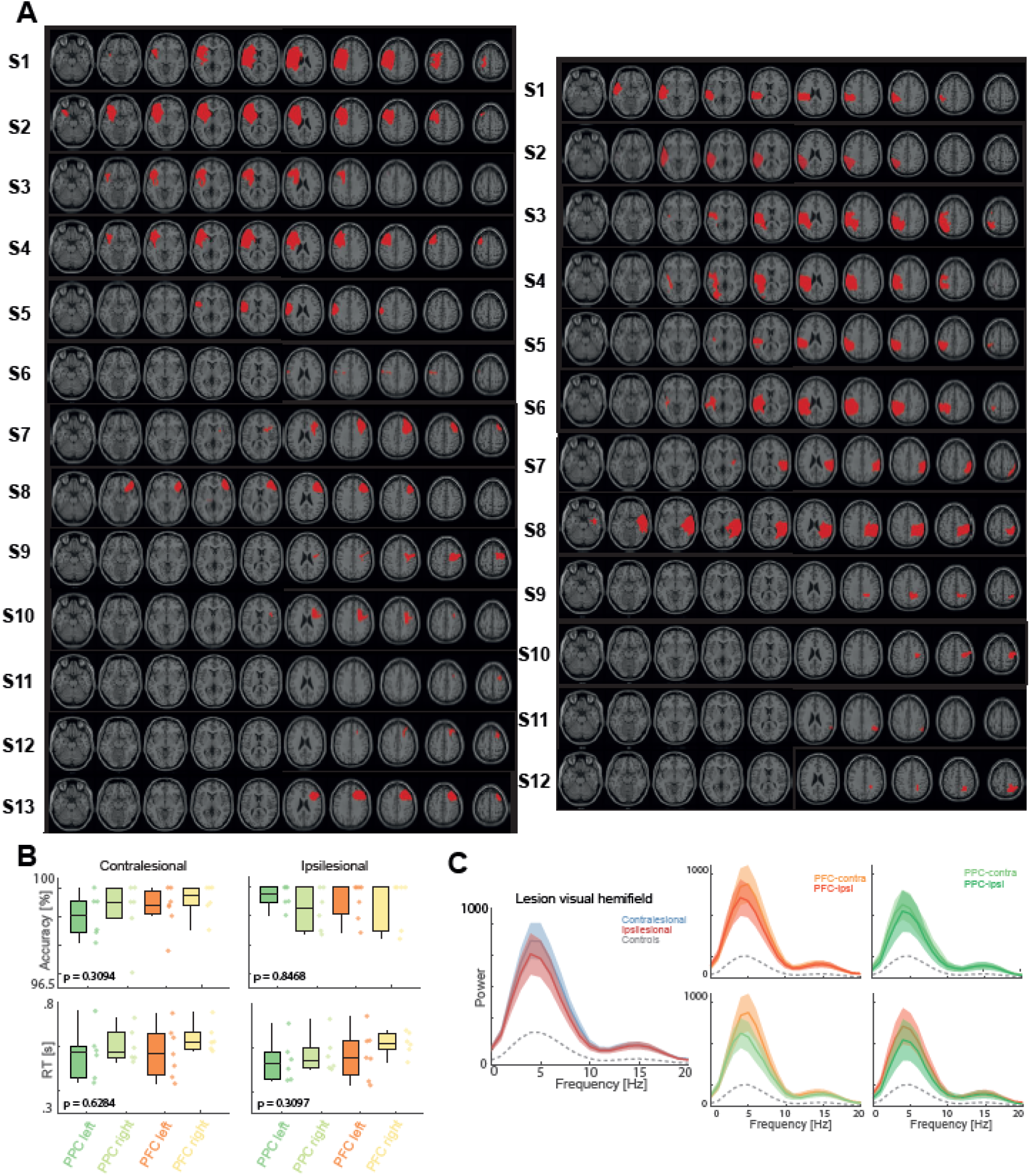
Lesion reconstruction and power spectra depending on the lesioned hemifield. (A) Reconstruction of the lesion of every patient with PFC (left) and PCtx (right) lesion. PFC lesions were in the left hemisphere of six patients and right hemisphere of seven patients. PCtx lesions were in the left hemisphere of six patients and right hemisphere of six patients. (B) Mean accuracy and reaction times for targets presented in the contra- or ipsilesional hemifield per group (whiskers indicate maximum and minimum data points not considered outliers; data points correspond to participants). (C) Group-level power spectra of the behavioral time courses (1/f-corrected) of the contralesional and ipsilesional hemifield. No difference was detected in patients when the target appeared on the contra- vs ipsilesional hemifield (no significant cluster, all uncorrected p > 0.2655). (C) Visualization for PFC and PCtx lesions separately. No significant difference was observed within nor between groups (p > 0.05).

**Figure S2 related to Figure 2.**
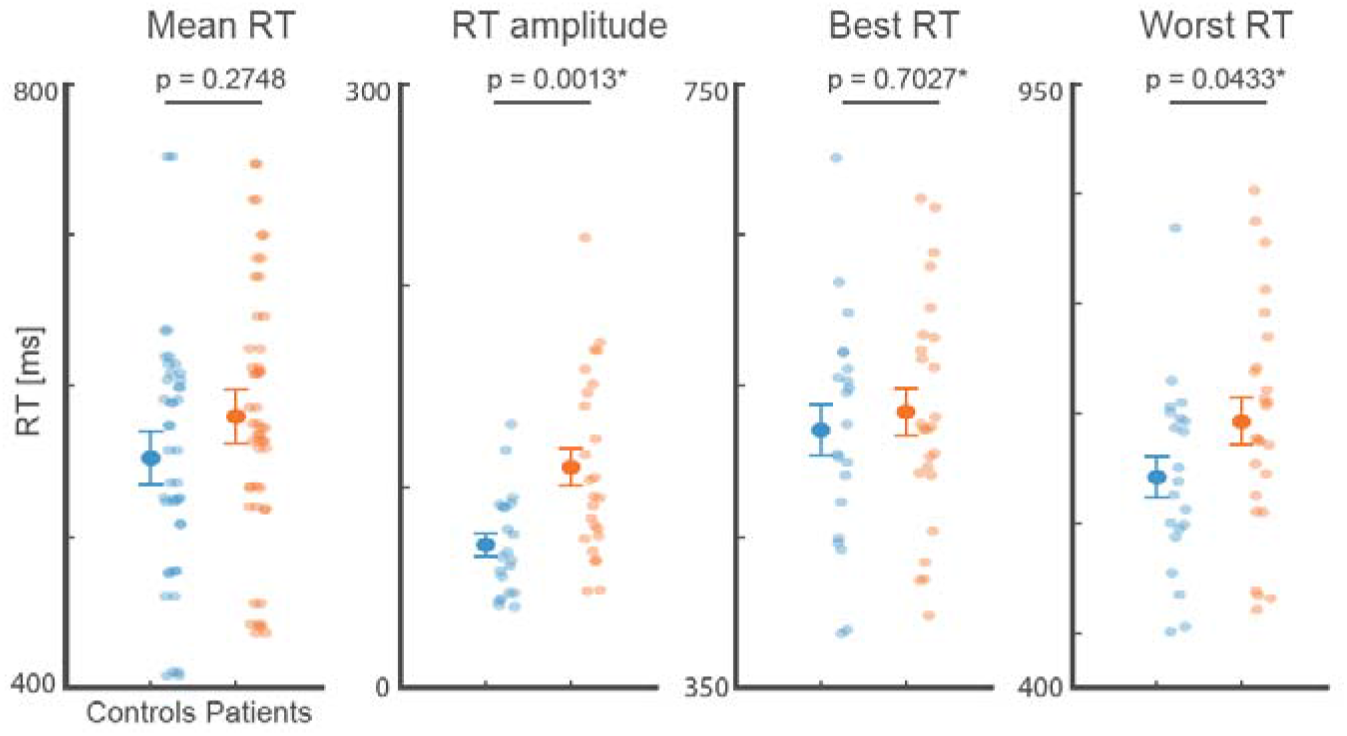
Band-pass filtered behavioral rhythmic sampling model results. Patients exhibited no significant difference in mean or best RT, but RT amplitude and worst RT were significantly increased (p = 0.0013), and worst RT (p = 0.0433) in the band-pass filtered data (0.25 – 7.75 Hz) as in the raw data.

**Figure S3 related to Figure 3C.**
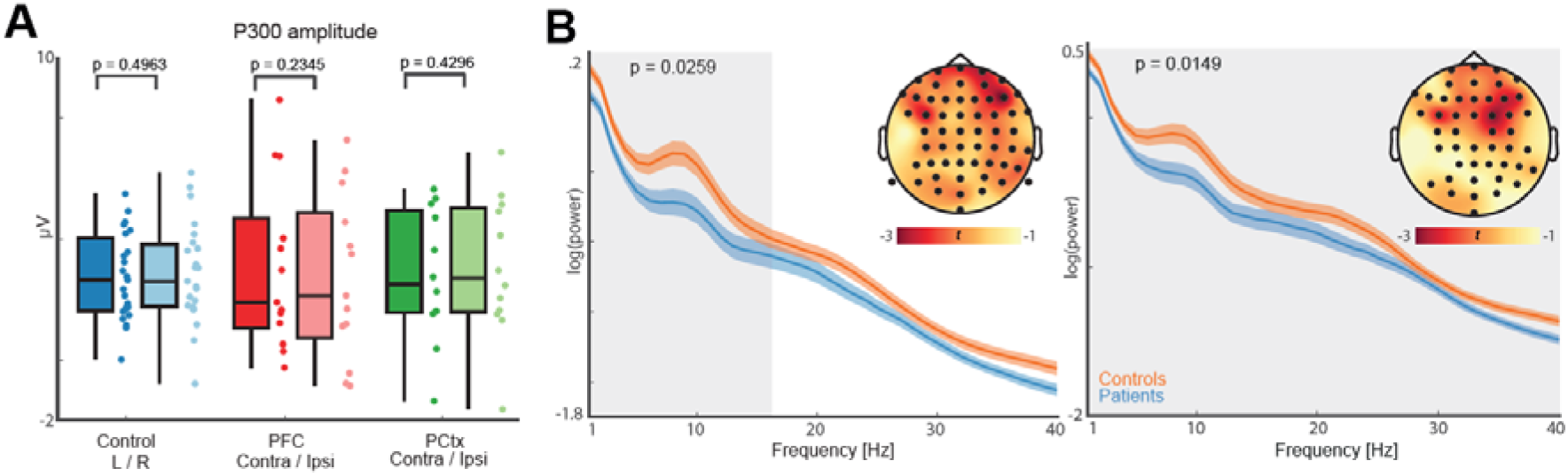
Unipolar averaging did not affect increased lower frequency oscillations. (A) Average P300 ERP peak in the time window of 200 to 350ms. No difference was observed between any of the groups. (B) Left: Grand-average power spectra of channels within the significant cluster (mean ± SEM; black dots indicate significant channels). Patients exhibited increased low frequency power in comparison to controls (1-16 Hz; p_cluster_ = 0.0259; grey shaded area). Right: Grandaverage power spectra of unipolar referencing to Cz. Patients showed increased oscillatory low frequency (1 – 40 Hz) power in comparison to controls (p_cluster_ = 0.0149; grey shaded area).

**Figure S4 related to Figure 4E and 4F.**
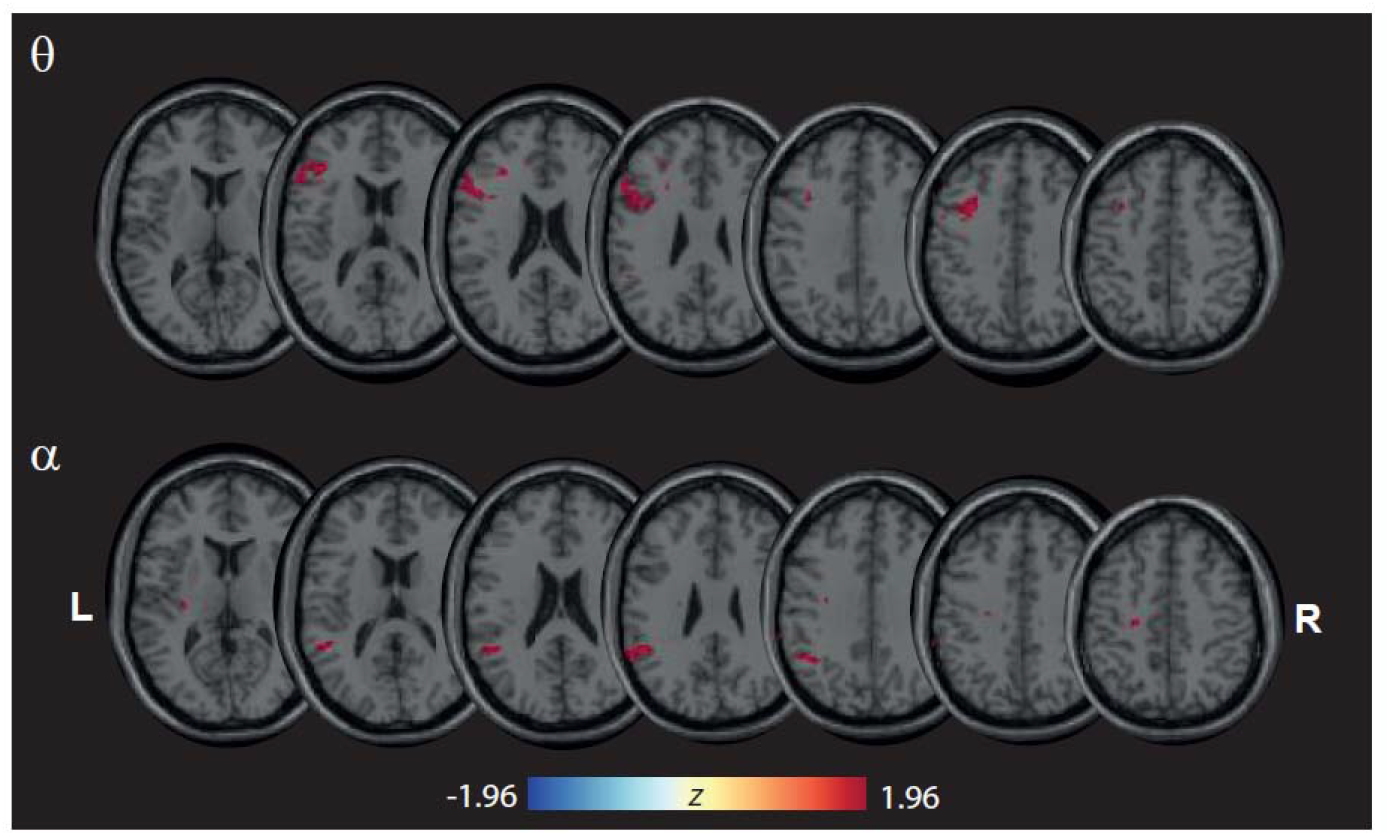
Lesions in the PFC and PCtx modulate different frequency bands. Spectrally-resolved VLSM maps showing *z*-values of only the lesion voxels with a highly significant effect on behavior power increase (FDR corrected p < 0.05). Lesions within the lateral prefrontal cortex predicted a behavioral theta power increase, whereas injury to the temporoparietal junction predicted a behavioral alpha power increase.

